# Transcriptional response to West Nile virus infection in the zebra finch (*Taeniopygia guttata*), a songbird model for immune function

**DOI:** 10.1101/103564

**Authors:** Daniel J Newhouse, Erik K Hofmeister, Christopher N Balakrishnan

## Abstract

West Nile Virus (WNV) is the one of most widespread arboviruses worldwide. WNV exists in a bird-mosquito transmission cycle where passerine birds act as the primary reservoir host. As a public health concern, the mammalian immune response to WNV has been studied in detail. Little, however, is known about the avian immune response to WNV. Avian taxa show variable susceptibility to WNV and what drives this variation is unknown. Thus, to study the immune response to WNV in birds, we experimentally infected captive zebra finches (*Taeniopygia guttata*). Zebra finches provide a useful model, as like many natural avian hosts they are moderately susceptible to WNV and thus provide sufficient viremia to infect mosquitoes. We performed splenic RNAseq during peak viremia to provide an overview of the transcriptional response. In general, we find strong parallels with the mammalian immune response to WNV, including up-regulation of five genes in the Rig-I-like receptor signaling pathway, and offer insights into avian specific responses. Together with complementary immunological assays, we provide a model of the avian immune response to WNV and set the stage for future comparative studies among variably susceptible populations and species.

## Introduction

West Nile virus (WNV) is a single-stranded RNA flavivirus that exists in an avian-mosquito transmission cycle, where birds (typically Passeriformes) act as the primary amplification hosts. In addition to birds, nearly 30 other non-avian vertebrate species have been documented as hosts (1). Although many WNV-infected hosts are asymptomatic, WNV infection can cause severe meningitis or encephalitis in those that are highly susceptible. Avian species for the most part exhibit low to moderate susceptibility. That is, individuals become infected and develop sufficient viremia for transmission via mosquito blood meal, but the hosts recover and avoid significant mortality, reviewed in (2). First described in 1937, WNV has not resulted in widespread avian decline throughout its historical range (3), perhaps due to host-parasite coevolution. However, the emergence of WNV in North America in 1999 has negatively impacted a wide range of populations (4,5). Surveys of North American wild birds have shown a variety of competent WNV hosts, with varying degrees of susceptibility, morbidity, and pathogenicity (2). American robins (*Turdus migratorius*) appear to be the main host in spreading WNV infection in North America (6), but infection appears most detrimental to members of Family Corvidae (7). Despite great variation in susceptibility, the mechanisms underlying this variation are primarily unknown (2).

Largely due to interest in human health implications, most work describing the host immune response to WNV infection has been performed in mammalian systems (8). From these studies, we know that in mammals, both the innate and adaptive arms are critical for virus detection and clearance (9, 10). Within the innate immune response, the retinoic acid-inducible gene 1(Rig-I)-like receptor (RLR) pathway appears to play a key role in viral clearance. This pathway recognizes viral products and initiates type I interferon expression (11). Mice lacking the viral recognition RLR genes in this pathway, DDx58 (Rig-I) and IFIH1 (MDA5), become highly susceptible to WNV infection (12). In the adaptive immune system, a broad range of components appear to play important roles in mounting a response, including antibody and CD4+ and CD8+ T cells (9,13,14).

Interestingly, major histocompatibility complex (MHC) class I genes are up-regulated post-infection (15,16). Viruses typically evade MHC class I detection (17,18), as MHC class I molecules bind and present viral peptides to CD8+ T cells. However, the purpose of WNV induced MHC expression is unclear.

While the mammalian immune response to WNV infection has been extensively studied, the avian immune response remains mostly unknown. Of the studies in birds, many involve experimentally infecting wild caught birds, reviewed in (2), or domestic chickens (*Gallus gallus*) (19). These studies primarily focus on viral detection, tissue tropism, antibody production, or lymphocyte counts (2,19, 20). Little is known about the molecular mechanisms driving the immune response to WNV infection, but see (21). Furthermore, current avian WNV studies suffer many challenges. Wild caught birds may be co-infected with other parasites (e.g. avian malaria) and are difficult to maintain in captivity for experimental infection studies. Chickens, although an avian model species, are highly resistant to WNV infection (22), uncommon hosts, and therefore are not ideal to describe the avian immune response to WNV infection. Passeriformes and Galliformes are also highly divergent bird lineages, with distinctive immune gene repertoires and architecture (23).

As passerine birds are the main hosts for WNV, we have sought to develop a passerine model to study the impacts of WNV infection on a taxonomically appropriate host (24). We have recently shown that zebra finches, *Taeniopygia guttata,* are moderately susceptible hosts for WNV (25). That is, WNV rapidly disseminates to avariety of tissues and is detectable in most samples by four days post-inoculation (dpi). Despite rapid development of sufficient viremia for arthropod transmission, zebra finches develop anti-WNV anitbodies, clear WNV by 14dpi, and avoid significant mortality (25). This moderate disease susceptibility is similar to what is observed in many natural WNV hosts. Zebra finches are also an established biomedical model system with a suite of genetic and genomic tools available (26).

In this study, we experimentally infected zebra finches and performed splenic RNAseq to describe their transcriptional response over the time course of infection. In doing so, we characterize the zebra finch immune response to WNV infection, explore expression of the avian RLR pathway in response to WNV, gain insights into the avian immune response to this widespread infectious disease, and uncover conserved evolutionary responses in avian and mammalian systems.

## Results

### Experimental infection

We challenged six individuals with with 10^5^ plaque forming units (PFU) WNV and sequenced RNA (Illumina RNAseq) isolated from spleens, an organ critical to the avian immune response. Three birds served as procedural controls and on day 0 were injected subcutaneously with 100 µL of BA1 media, as previously described (27). Peak viremia occurrs at 4.6 ±1.7 dpi as quantified via RT-PCR (25) and thus, we characterized the transcriptional response leading to (2dpi, n=3) and at peak viral load (4dpi, n=3) in the present study. WNV RNA was detected by culture in lung and kidney RNA pools of 2 out of 3 birds sampled at day 2, and all 3 birds sampled at 4dpi. These findings were verified by semi-quantitative RT-PCR. Because WNV is rarely detected in spleen by 2dpi, but all birds previously inoculated at 10^5^ PFU developed WNV antibodies [25] we treated all six birds inoculated with WNV as being infected.

### Sequencing results & read mapping

We obtained 18-30 million paired-end, 100bp reads for each sample and removed 0.57-1.24% of the total bases after adapter trimming (Supplementary Table S1). On average, 79.0-80.8% total trimmed reads mapped to the zebra finch reference genome (Supplementary Table S2), corresponding to 18,618 Ensembl gene IDs. Of these, 14,114 genes averaged at least five mapped reads across all samples and were utilized for differential expression (DE) analyses.

### Sample clustering & differential expression

We tested for DE two ways: as pairwise comparisons between treatments to identify specific genes with *DEseq2* (28) and as a time-course grouping genes into expression paths with *EBSeqHMM* (29). To visualize patterns of expression variation among samples, we conducted principal component analysis (PCA) and distance-based clustering (Supplemental Figures S1 & S2). The first three principal components explained 93.04% of the variance in gene expression, but none of the PCs were significantly correlated with treatment (ANOVA, PC1: p = 0.288, PC2: p = 0.956, PC3: p = 0.202). Although this finding suggests that much of the expression variation was independent of the experimental treatment, pairwise comparisons revealed genes that were DE between treatments.

When comparing Control vs. 2dpi, we found 161 differentially expressed genes (adjusted p < 0.10, average log_2_ fold-change (FC) = 1.74). This gene list includes several immune related genes associated with the innate (e.g. IL18) and adaptive (e.g. MHC IIB) immune system (Table 1, Figure 1). Sixty-five genes were differentially expressed between Control and 4dpi (average log_2_FC = 1.61), also with several immune relevant genes including five genes in the RLR pathway (Table 1, Figure 2, Figure 3). Lastly, we observed 44 DE genes between 2dpi vs. 4dpi individuals (average log_2_FC = 1.56). Three of these have described functions in immunity. The complete list of *DEseq2* DE genes (adjusted p < 0.10) across all comparisons can be found in Supplementary Table S3 and immune relevant DE genes are listed in Table 1 along with their gene name, log_2_ fold change, padj value, and comparison to mammalian studies. We also combined 2dpi and 4dpi cohorts and compared with control, but due to high variation in gene expression between days 2 and 4 dpi, we only found 16 DE genes (average log_2_FC = 1.64) between Control and Infected cohorts, one of which was associated with immunity.

**Table 1.** Candidate immune genes differentially expressed in the present study and comparisons with mammals.

| Ensembl ID | Gene Name | Log<sub>2</sub> Fold Change | padj | Regulation Pattern Observed | Regulation Pattern in Mammals | Reference |
| --- | --- | --- | --- | --- | --- | --- |
| ***Control vs Infected*** |  |  |  |  |  |  |
| ENSTGUG00000013615 | NFKBIZ | 0.73 | 0.064 | Up | Up | 45 |
| ***Control vs 2dpi*** |  |  |  |  |  |  |
| ENSTGUG00000000297 | IL18 | 1.01 | 0.010 | Up | No change | 51 |
| ENSTGUG00000000678 | TIM1 | 1.49 | 7.99E-05 | Up | Up | 41,42 |
| ENSTGUG00000001485 | IRF6 | -2.09 | 0.037 | Down | Up | 43 |
| ENSTGUG00000003354 | NKRF | -2.35 | 4.85E-05 | Down | Unknown |  |
| ENSTGUG00000005295 | C-C motif chemokine | 2.08 | 0.007 | Up | Up | 43,44,45 |
| ENSTGUG00000008638 | UCHL1 | -1.91 | 0.029 | Down | Unknown |  |
| ENSTGUG00000008991 | APOD | 1.76 | 0.053 | Up | Up | 46 |
| ENSTGUG00000009454 | IFITM10 | 1.24 | 2.71E-04 | Down | Unknown |  |
| ENSTGUG00000009769 | TNFRSF13C | 0.89 | 0.010 | Up | Unknown |  |
| ENSTGUG00000015634 | Novel gene (MHC IIB) | 2.23 | 0.001 | Up | Up | 48 |
| ENSTGUG00000016383 | SIGLEC1 | 1.39 | 0.046 | Up | Up | 45 |
| ENSTGUG00000017149 | Novel gene (MHC IIB) | 1.57 | 0.099 | Up | Up | 48 |
| ***Control vs 4dpi*** |  |  |  |  |  |  |
| ENSTGUG00000001516 | DDx58 | 1.50 | 1.45E-08 | Up | Up | 45 |
| ENSTGUG00000002144 | IRF4 | 1.39 | 0.022 | Up | Up | 43 |
| ENSTGUG00000002305 | LY86 | -1.07 | 1.70E-05 | Down | Unknown |  |
| ENSTGUG00000002516 | DHx58 | 1.67 | 4.05E-06 | Up | Up | 45 |
| ENSTGUG00000004105 | ADAR | 1.13 | 1.70E-05 | Up | Up | 47 |
| ENSTGUG00000006914 | IFIH1 | 0.95 | 0.093 | Up | Up | 45 |
| ENSTGUG00000007454 | TNFRSF13B | 1.61 | 0.010 | Up | Unknown |  |
| ENSTGUG00000008354 | IFIT5 | 2.97 | 1.07E-09 | Up | Unknown |  |
| ENSTGUG00000008788 | EIF2AK2 | 2.15 | 9.86E-07 | Up | Up | 43,69 |
| ENSTGUG00000009162 | PXK | 1.19 | 0.011 | Up | Unknown |  |
| ENSTGUG00000009536 | TRIM25 | 1.24 | 0.001 | Up | Up | 45 |
| ENSTGUG00000009838 | IRF7 | 0.80 | 0.047 | Up | Up | 45 |
| ENSTGUG00000011784 | ZC3HAV1 | 1.31 | 7.09E-08 | Up | Up | 45 |
| ENSTGUG00000017534 | MOV10 | 1.29 | 0.017 | Up | Up | 47 |
| ***2dpi vs 4dpi*** |  |  |  |  |  |  |
| ENSTGUG00000003354 | NKRF | 1.70 | 0.033 | Up | Unknown |  |
| ENSTGUG00000005206 | ADA | -1.48 | 0.001 | Down | Unknown |  |
| ENSTGUG00000011784 | ZC3HAV1 | 0.82 | 0.058 | Up | Up | 45 |

**Figure 1.**
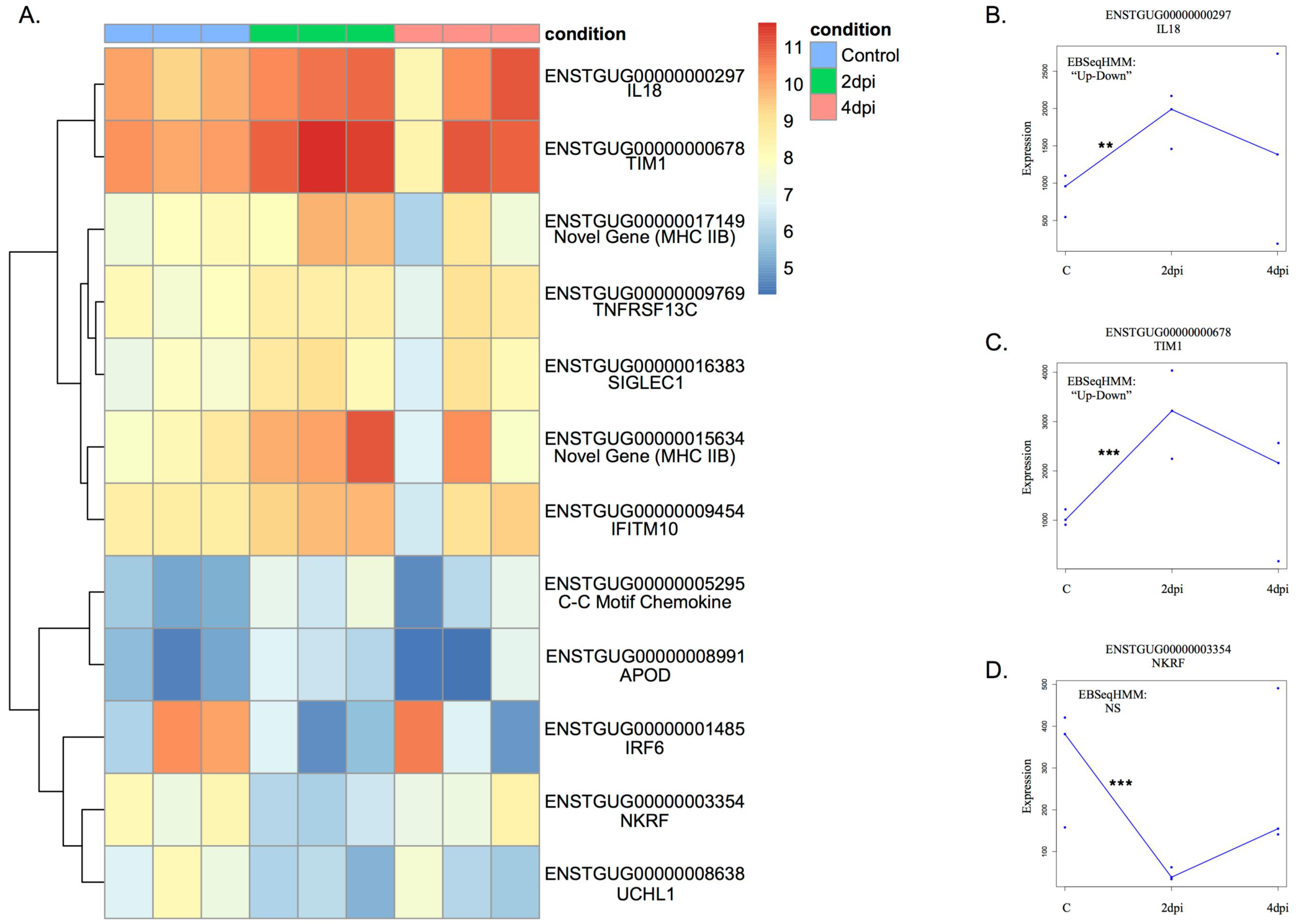
Immune genes differentially expressed between day 2 post-inoculation and control. A) Heatmap of expression levels (log transformed read counts) across alltreatments of immune genes differentially expressed at 2dpi relative to control. B-D) Expression values (normalized read counts) for three key immune genes and their regulation pattern classification by *EBSeqHMM*. Asterisks represent statistical significance in *DEseq2* analysis after FDR correction (* p<0.10, ** p<0.05, *** p<0.01).

**Figure 2.**
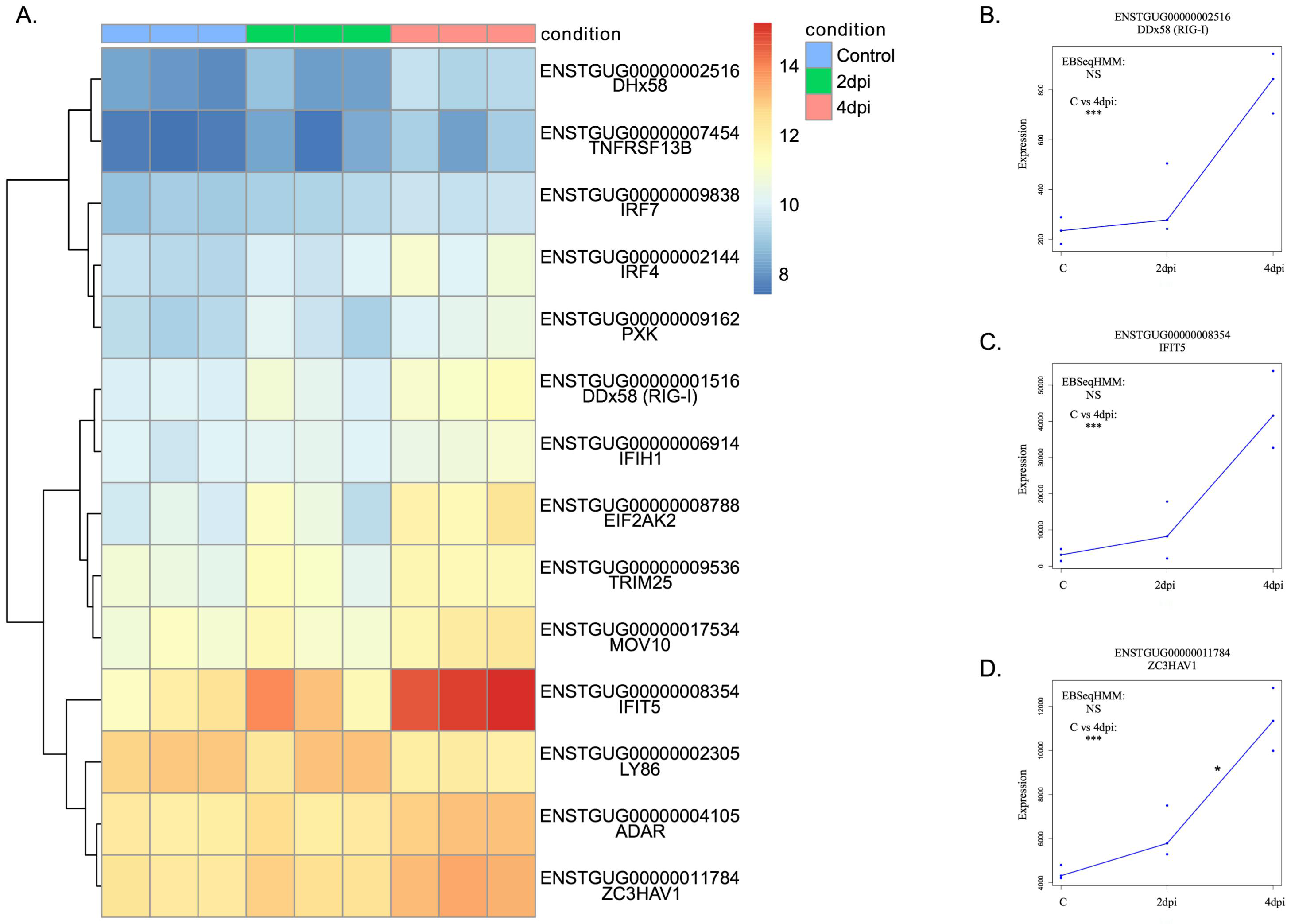
Immune genes differentially expressed between day 4 post-inoculation and control. A) Heatmap of expression levels (log transformed read counts) across alltreatments of immune genes differentially expressed at 4dpi relative to control. B-D) Expression values (normalized read counts) for three key immune genes and their regulation pattern classification by *EBSeqHMM*. Asterisks represent statistical significance in *DEseq2* analysis after FDR correction (* p<0.10, ** p<0.05, *** p<0.01).

**Figure 3.**
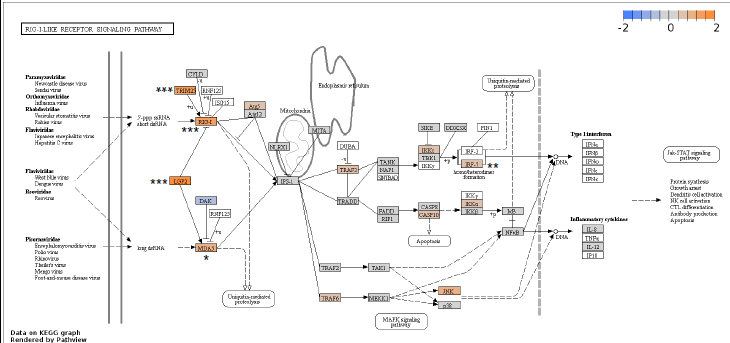
Regulation of the zebra finch RLR pathway. Color represents log_2_foldchange between Control and 4dpi. Asterisks represent statistical significance in *DEseq2* analysis after FDR correction (* p<0.10, ** p<0.05, *** p<0.01).

When analyzed for DE as a time course in *EBSeqHMM*, 686 genes showed evidence of differential expression (posterior probability > 0.99, FDR < 0.01). Most DE genes (n = 561) were suppressed following infection (“Down-Down”) or varied between conditions. Seventy-five genes were “Up-Down”, 49 were “Down-Up” and 1 was “Up-Up”. As expected, we found overlap of several immune genes between the two analyses. For example, IL18, APOD and IFITM10 are “Up-Down" (Supplementary Table S5) and this trend is reflected in the *DEseq2* Control vs 2dpi analysis (Figure 1).

### Functional annotation of differentially expressed genes

To place differentially expressed genes into groups based on their biological function, we performed a gene ontology (GO) analysis using the *GOfinch* tool (30). As above, we conducted GO analyses based on multiple pairwise analyses of gene expression. The strongest evidence of functional enrichment was observed in the contrast of Control vs. 4dpi. This list of 65 differentially expressed genes showed functional enrichment of 55 GO terms (Fisher’s Exact Test, FDR p < 0.05 (Benjamini Hochberg method [31])). It is in this contrast where the immune response manifests itself most strongly, with a large list of immune-related categories (Supplementary Table S4) and a broad range of immune functioning genes differentially expressed (n=14, Tables 1 & 2). When comparing genes differentially expressed between Control and 2dpi, only one GO category was strongly enriched (“integral to membrane”, expected = 15, observed = 35, p = 0.00065). In this contrast we also observed several slightly enriched immune related GO categories such as “inflammatory response” and “negative regulation of T cell migration” but these fall outside of our significance threshold (Supplementary Table S4) and reflect only few genes (e.g. APOD and IL18) that were DE in the Control vs. 2dpi analysis. A set of 57 enriched GO categories describe genes DE between 2 and 4dpi. Although some of these involved T and B cell regulation, these categories were also only represented by a single gene each (ADA) (Table 2, Supplementary Table S4). We found no evidence of functional enrichment among DE genes between Control vs. Infected (2dpi and 4dpi) cohorts.

**Table 2.**
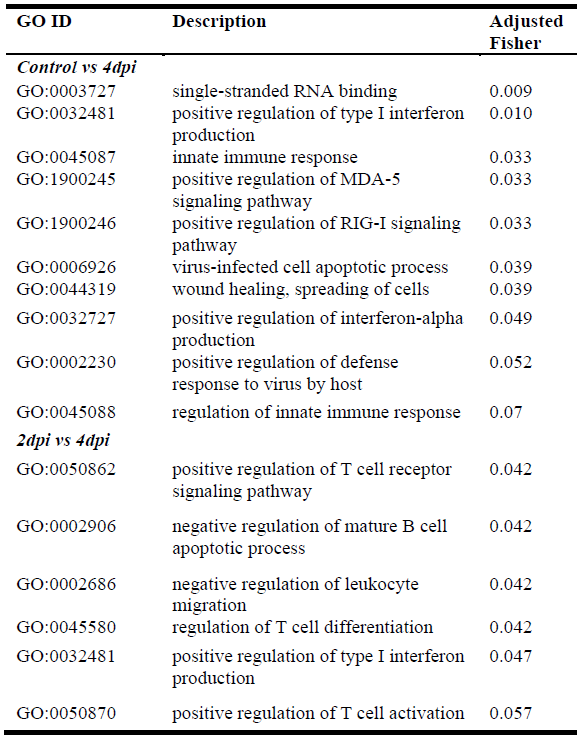
Select immune related gene ontology (GO) categories among DEseq2 significantly differentially expressed genes (adjusted p value < 0.10).

| GO ID | Description | Adjusted Fisher |
| --- | --- | --- |
| *Control vs 4dpi* |  |  |
| GO:0003727 | single-stranded RNA binding | 0.009 |
| GO:0032481 | positive regulation of type I interferon production | 0.010 |
| GO:0045087 | innate immune response | 0.033 |
| GO:1900245 | positive regulation of MDA-5 signaling pathway | 0.033 |
| GO:1900246 | positive regulation of RIG-I signaling pathway | 0.033 |
| GO:0006926 | virus-infected cell apoptotic process | 0.039 |
| GO:0044319 | wound healing, spreading of cells | 0.039 |
| GO:0032727 | positive regulation of interferon-alpha production | 0.049 |
| GO:0002230 | positive regulation of defense response to virus by host | 0.052 |
| GO:0045088 | regulation of innate immune response | 0.07 |
| *2dpi vs 4dpi* |  |  |
| GO:0050862 | positive regulation of T cell receptor signaling pathway | 0.042 |
| GO:0002906 | negative regulation of mature B cell apoptotic process | 0.042 |
| GO:0002686 | negative regulation of leukocyte migration | 0.042 |
| GO:0045580 | regulation of T cell differentiation | 0.042 |
| GO:0032481 | positive regulation of type I interferon production | 0.047 |
| GO:0050870 | positive regulation of T cell activation | 0.057 |

We also conducted a similar analysis of genes identified as DE by *EBseqHMM*, which revealed two (Up-Up), 107 (Up-Down), 12 (Down-Up) and 73 (Down-Down) significantly enriched GO categories (adjusted p < 0.05, Supplementary Table S5). Interestingly, Up-Down GO categories had the strongest representation of immune related GO terms, including “cytokine receptor activity” (expected = 0, observed = 3 p = 0.015) and “positive regulation of interferon-gamma production” (expected = 0, observed = 2, p = 0.028). However, we also observed a strong immune signature among Down-Down Genes (GO Term “Immune Response”, expected = 3, observed = 11, p = 0.013), which represents several innate immune genes including three complement genes and interferon gamma. Twelve categories were significantly enriched among “Down-Up” genes, including “double-stranded RNA binding” (expected = 0, observed = 2, adjusted p = 0.049). One of the observed genes in this category is OASL, which has been demonstrated to have antiviral activity towards WNV in chickens (21). Lastly, as in the *DEseq2*-based analysis, we also detected a strong enrichment signature of membrane proteins. Genes annotated as “integral to membrane” were highly enriched among those showing an Up-Down pattern (expected = 6, observed =14, adjusted p < 0.028, Supplementary Table S5). Combined, we find broad overlap in GO representation between the *EBseqHMM* and *DEseq2* approaches.

In addition to placing genes into broad systematic functions in the GO analysis, we were also interested in placing our gene expression results in the context of immune pathways of interest. The RLR antiviral pathway is critical to WNV clearance in mammals (12) and appears important in mounting an immune response to avian influenza in ducks (32–34). Utilizing *Pathview v1.8.0* (35), we find that WNV infection induces the RLR pathway. Five genes, including the two RLR genes, DDx58 and IFIH1, which encode the Rig-I and MDA5 viral detection molecules, are significantly up-regulated (Table 1, Figure 3, Supplementary Figure S3). We detect expression of 36/37 genes in the pathway, many of which are also up-regulated, though not always significantly.

### De novo transcriptome analysis

To reveal any genes responding to infection missed by our genome based analysis, we created a reference transcriptome with *Trinity v2.1.1* (35) and annotated against the NCBI non-redundant database with *DIAMOND*, a *BLASTx*-like aligner (37). Our *de novo* transcriptome assembly resulted in 393,408 *Trinity* transcripts. We visualized our annotated transcriptome with *MEGAN* (38), which places genes into KEGG pathways. In doing so, we confirmed many expression results from the genome-based approach and importantly, this analysis revealed expression of two transcripts with homology to interferon alpha, which was absent (but annotated) in the genome based analysis described above. Combined, our genome-based and reference-free approaches detect regulation of the complete RLR pathway in zebra finches.

## Discussion

We have characterized the zebra finch transcriptional response to WNV infection. Overall, we find that as in mammalian systems, components of both the adaptive and innate immune pathways are activated following infection. While WNV is primarily an avian specific infectious disease, most work describing the host immune response to infection has been performed in mammals. Despite genomic, physiological and evolutionary differences between birds and mammals, the host immune response shows broad similarity between taxa (Table 1).

We were particularly interested in the role of the innate RLR pathway. This pathway mounts an antiviral innate immune response and is critical for WNV detection and clearance in mammals (12). By combining genome and transcriptome-based approaches, we show here that the RLR pathway in zebra finches is induced by WNV infection, as gene expression is detected throughout the pathway. Furthermore, five genes in this pathway are significantly up-regulated at 4dpi (Figure 3, Supplementary Figure S3), including IFIH1 and DDx58 (Figure 2B), which encode molecules that recognize WNV particles in mammals (39). This results in a corresponding over-representation of genes in the RLR pathway GO categories (Table 2). While no studies have investigated the role of the RLR following WNV infection in birds, this pathway appears important for avian influenza clearance in ducks (32-34), Buggy Creek virus clearance in house sparrows (39), and likely for the broad avian antiviral immune response, including WNV.

We observed other parallels with mammals as well (Table 1). For example, T-Cell Immunoglobulin Mucin Receptor 1 (TIM1) is up-regulated at 2dpi in zebra finches (Figure 1A, C). In human cell lines, expression of TIM1 promotes infection of WNV virus like particles (VLPs) (41, 42), suggesting that the up regulation of TIM1 seen in zebra finches may promote viral entry as well. Similarly, C-C motif chemokine (ENSTGUG00000005295) is up-regulated in our study at 2dpi and in previous human cell line and mouse experiments, suggesting a conserved role in chemokine production following WNV infection (43–45). Apolipoprotein D (APOD), a gene typically involved in brain injury and potentially responding to the neurodegenerative nature of WNV, is up-regulated in WNV infected mice (46), as well as in our study. Two interferon stimulated genes (ISGs), ADAR and MOV10 are both significantly up-regulated at 4dpi relative to control. Schoggins et al. (47) showed ADAR expression to enhance WNV replication and MOV10 expression to have antiviral activity. While further testing of these genes is needed to validate their roles in avian WNV infection, they nonetheless offer insights into a broad range of conserved responses between mammals and birds.

Within the adaptive immune response, the role of the MHC in the host response to WNV is also particularly interesting. The MHC plays a key role in antigen processing and presentation. The MHC comprises two main classes (Class I & II) and both are up-regulated in mammals following WNV infection (48, 15, 16). Similarly, two genes encoding MHC class IIB proteins are significantly up-regulated in zebra finches at 2dpi (Figure 1). Unlike mammals, however, we found that MHC class I is not significantly DE in any comparison. In mammals, upregulation of MHCI may not be adaptive for the host, as upregulation may actually be a mechanism by which the virus evades Natural Killer (NK) cell detection by the innate immune system (15). It has also been suggested that MHC up-regulation is a byproduct of flavirus assembly (49). Interestingly, at 2dpi, interleukin-18 (IL18) is significantly up-regulated (Table 1, Figure1 A, B). IL18 can enhance NK cell activity (50) and is potentially a mechanism by which the immune system can counteract WNV evasion strategies via NK cell activation, although further testing is needed to quantify NK cell activity in zebra finches to support this hypothesis.

Despite many similarities, several immune genes differentially expressed in our analyses have not been previously reported in the mammalian WNV literature or are expressed differently in zebra finches (Table 1). For example, at 2dpi, the proinflammatory cytokine IL18 was significantly up-regulated in zebra finches (Figure 1), contrasting a previous study in human cell lines, which show no difference in IL18 expression following WNV infection (50). Furthermore, interferon regulatory factor 6 (IRF6) was down-regulated at 2dpi, but up-regulated in human macrophages following infection (43). Another significantly down-regulated gene at 2dpi, ubiquitin carboxyl-terminal hydrolase L1 (UCHL1), has been shown to suppress cellular innate immunity in human cell lines infected with high-risk human papilloma virus (52). Down-regulation restores functional pattern recognition receptor (PRR) pathways (e.g. RLR). The down-regulation of UCHL1 here thus is associated with the previously described regulation of the PRR RLR pathway in this study (Table 1). Lastly, interferon-induced protein with tetratricopeptide repeats (IFIT) and interferon-inducible transmembrane proteins (IFITM) gene families are known innate antiviral proteins and have been shown to restrict WNV entry in human cells lines (47, 53). Both IFIT5 and IFITM10 are up-regulated (Figure 2A, C) in our study and yet, to our knowledge, neither have previously been implicated in the host immune response to WNV. This potentially reveals an avian specific function of IFIT5 and IFITM10.

Like many passerine birds infected in nature, zebra finches are moderately susceptible to WNV, developing sufficient viremia to serve as competent hosts, but generally resisting mortality due to infection (25). While there are clear differences among treatments in terms of differentially expressed genes (Table 1), the weak effect of treatment on overall expression profile (Supplemental Figure S1 & S2). may be a reflection of this moderate susceptibility. Most zebra finches are able to clear WNV inflection by 14 dpi (25). In pairwise comparisons, there are between 16 and 161 differentially expressed genes, depending on the treatment comparison (Supplementary Table S3). WNV infection intensity varies among tissues (20), but due to the spleen’s important role in the avian immune system (54,55), we expect these results to be representative of the overall immune response.

Functional enrichment of several immune GO terms primarily appears in Up-Down path defined by *EBseqHMM* (Supplementary Table S5), as many genes in the immune system are up-regulated post-infection, as also seen in the *DEseq2* analysis (Table 1, Figure 1, Figure 2). In both the *EBseqHMM* and *DEseq2* analyses, most of the significant immune GO categories are innate immune responses, although adaptive immune categories involved in B/T cell proliferation appear in both (Table 2, Supplementary Tables S4 & S5). Similar to the mammalian model, broad organismal processes, encompassing the entire immune response, are represented in the zebra finch response to WNV.

The zebra finch was the second bird to have its genome sequenced (26). However, the zebra finch genome assembly and gene annotation remain incomplete. Immune genes in particular are difficult to reconstruct in genomes due to their complex evolutionary history (e.g. MHC, [23]). Our mapping results indicated that roughly 80% of our reads mapped to the genome. While the unmapped 20% could contain poor quality reads or non-avian sequences, it also represents zebra finch reads that could not be appropriately mapped to the reference. Thus, we took a genome-independent approach and assembled a *de novo* zebra finch transcriptome from our nine paired-end samples. Interestingly, RLR induced interferon expression was not detected in the genome-based analysis. The transcriptome analysis, however, revealed expression of two transcripts with homology to interferon alpha, thus completing the pathway from virus detection to interferon production.

We have begun to develop the zebra finch as an avian model for the host response to WNV infection. We show here that the zebra finch immune response is largely conserved with that seen in mammalian-based studies (Table 1). Additionally, we identify many components of the immune system that have not been previously implicated in the host immune response to WNV. This potentially reveals an avian-specific immune response and highlights avenues for future research. Combined with our recent immunological characterization (25), we have broadly described the immuneresponse of a moderately susceptible avian host for WNV. This sets the stage for future comparative work to uncover the genetic basis of variable avian susceptibility to WNV infection.

## Methods

### Experimental Setup

All animal use was approved by the USGS National Wildlife Health Center Institutional Animal Care and Use Committee (IACUC Protocol: EP120521) and this study was performed in accordance with USGS IACUC guidelines. The experimental infection setup is described in detail in (25). Briefly, nine female zebra finches were randomly divided into three cohorts, one unchallenged and two challenged (n = 3 each). Birds were challenged subcutaneously with 100ul BA1 media containing 10^5^ plaque-forming units (PFU) of the 1999 American crow isolate of WNV (NWHC 16399-3) and sacrificed at 2 and 4 dpi, corresponding to peak viremia. Uninfected individuals were injected with 100ul BA1 media and sacrificed at 4dpi. WNV infection was confirmed by RT-PCR, as previously described (26), in lung and kidney pooled tissue (25). Spleens from each individual were removed, placed into RNAlater (Qiagen, Valencia, CA USA), and frozen at −80 °C until RNA extraction.

### RNA extraction & sequencing

Whole spleen tissue was homogenized in Tri-Reagant (Molecular Research Company) and total RNA was purified with a Qiagen RNeasy (Valencia, CA USA) mini kit following the manufacturer’s protocol. RNA was DNAse treated and purified. Purified RNA was quality assessed on a Bioanalyzer (Agilent, Wilmington, DE USA) to ensure RNA quality before sequencing (RIN = 6.6-8.1). All library prep and sequencing was performed at the University of Illinois Roy J. Carver Biotechnology Center. A library for each sample was prepared with an Illumina TruSeq Stranded RNA sample prep kit. All libraries were pooled, quantitated by qPCR, and sequenced on one lane of an Illumina HiSeq 2000 with a TruSeq SBS Sequencing Kit producing paired-end 100nt reads. Reads were analyzed with Casava 1.8.2 following manufacturer’s instructions (Illumina, San Diego, CA). Sequencing data from this study have been deposited in the NCBI Sequence Read Archive (BioProject: PRJNA352507).

### Adapter trimming & read mapping

We removed Illumina adapters from reads with *Trim Galore! v0.3.7* (http://www.bioinformatics.babraham.ac.uk/projects/trim_galore/) which makes use of *Cutadapt v1.7.1* (56). Reads were then mapped to the zebra finch genome (*v3.2.74*,6) using *TopHat v2.0.13* (57), which utilizes the aligner *Bowtie v2.2.4* (58). We specified the library type as fr-firststrand in *TopHat2*. Successfully mapped reads were converted from SAM to BAM format with *SAMtools View v1.2* (59, 60) and counted in *htseq-countv0.6.0* specifying ‘-s rev’ (61). This assigned zebra finch Ensembl gene IDs and we only retained genes that mapped an average of five times across each sample.

### Differential expression

Gene counts were then normalized for read-depth and analyzed for DE in *DEseq2v1.8.1* (28). We analyzed DE across four comparisons: Control vs. Infected, Control vs. 2dpi, Control vs. 4dpi, and 2dpi vs. 4dpi. We visualized expression profiles in *R v3.3.0* (62) by PCA with the R package pcaExplorer (63), and hierarchical clustering heat maps with the ggplot2 library (64) following the *DEseq2* manual. *DEseq2* tests for DE with a Wald test and genes were considered differentially expressed if the Benjamini & Hochberg (31) false discovery rate (FDR) correction for multiple testing p value < 0.10. We chose this significance threshold as *DEseq2* is generally conservative in classifying DE (65). Furthermore, this cutoff has been used in other RNAseq experimental infection studies (66). We plotted genes of interest individually with the plotCounts function in *DEseq2* and clustered expression profiles of these genes with the pheatmap R library to view expression levels across samples and treatments.

We tested DE genes for under and over representation of gene ontology (GO) categories with the *CORNA* software (http://www.ark-genomics.org/tools/GOfinch, [30]). Significant DE genes were tested against all genes in our dataset. Statistical significance was determined using Fisher’s exact tests corrected for multiple hypothesis testing (p < 0.05). To visualize DE results in the context of the RLR pathway, we utilized *Pathviewv1.8.0* (35) to plot the log fold change of each gene detected in our dataset into the Kyoto Encyclopedia of Genes and Genomes (KEGG) pathway (KEGG ID = 04622) (67, 68).

### Time-course gene expression

In addition to the pair-wise comparisons performed in *DEseq2*, we were interested in understanding how clusters of genes are differentially expressed over the time course of infection. Thus, we performed DE analyses in *EBSeqHMM* (29). *EBSeqHMM* utilizes a bayesian approach with a hidden Markov model to identify DE between ordered conditions. Genes are then grouped into expression paths (i.e. “Up-Down”, “Down-Down”), in which DE occurs when expression paths change between at least one adjacent condition. For example, a gene up-regulated at both 2dpi relative to control and 4dpi relative to 2dpi would be classified as “Up-Up”. We included three time points, with control individuals classified as t1, 2dpi as t2 and 4dpi as t3. Genes were considered DE at posterior probability > 0.99 and FDR < 0.01. We chose a more stringent cutoff in this analysis as *EBseq* can be liberal in classifying differential expression (65) and based on visual inspection of expression profiles. Using genes identified as DE we performed the GO analysis described above.

### De novo transcriptome analysis

The use of Ensembl gene annotations for read counting restricts analyses to previously annotated genes. To test for potential regulation in any unannotated genes, we created a *de novo* transcriptome from our trimmed, paired-end reads using *Trinity v2.0.6* (36). We used default Trinity parameters with the exception of server specific settings (e.g.-max_memory 100G) and strand specificity (-SS_lib_type RF). To annotate our transcriptome, we used the *BLASTX*-like aligner *DIAMOND v0.7.11* (37), which utilizes NCBI’s non redundant database. We visualized results in *MEGAN v5.10.6* (38), which places transcripts into functional pathways (e.g. KEGG), facilitating comparisons to our genome based analysis.

## Acknowledgements

The authors wish to thank Melissa Lund for technical assistance. Funding provided by U. S. Geological Survey National Wildlife Health Center and East Carolina University. Any use of trade, firm, or product names is for descriptive purposes only and does not imply endorsement by the U. S. Government

## Additional Information

Sequencing data have been deposited in the Sequence Read Archive (SRA) under accession PRJNA352507. The authors declare no competing financial interests

